# *Priestia megaterium* ILBB592 based biofertilizer increases the efficiency of phosphorus fertilization, positively affects soybean nutrition and yield and modifies the rhizospheric bacterial community

**DOI:** 10.1101/2024.12.17.629018

**Authors:** Pablo Torres, Pablo Fresia, Elena Beyhaut, María José Cuitiño, Nora Altier, Eduardo Abreo

## Abstract

Phosphorus (P) is needed by soybean plants to form vital molecules like ATP and nucleotides and perform plant functions like photosynthesis and Nitrogen biological fixation at nodules. Because P concentration in the soil solution is naturally low, P fertilization is applied to satisfy soybean requirements in high yielding crops. However, a small portion of added P is readily available to the plants, while most is retained by the soil matrix and eventually made available or lost through erosion and run-off to water bodies, where it promotes eutrophication. Therefore, technologies that enhance P uptake by crops could lead to lower P inputs and outputs from soybean cropping systems, improving sustainability. Priestia megaterium ILBB592 and Bacillus pumilus ILBB44 are two PGPR with P mobilization features that were single formulated as biofertilizers and co-inoculated with Bradyrhizobium elkanii on seeds of soybean Genesis 5602 in two consecutive years in the field, with and without P fertilization. Co-inoculation with ILBB592 improved plant P uptake, along with N and K, with and without P fertilization. Diversity was increased by co-inoculation with ILBB44 and ILBB592, and predicted genes related with P cycling in the rhizospheric soils were also augmented after co-inoculation with ILBB592, mostly with P fertilization. Shoot dry weight and yield were also improved although effect on yield was not statistically significant. P fertilization alone had no effect in the first year but showed some effect on nodules dry weight, P uptake and yield on the second year, suggesting a sufficient P level was obtained after repeated fertilizations. ILBB592 biofertilizer showed a positive effect on most plant parameters, including nodulation and P uptake in both years and both P fertilization regimes and is therefore considered as a useful technology to reduce P fertilization without jeopardizing plant performance.

## Introduction

In soils, phosphorus (P) accumulates in organic and inorganic forms that are not readily available to plants, requiring the action of organic acids, phosphatases and phytases excreted by roots and microorganisms to render it bioavailable in the soil solution (Cheng et al., 2023). Since the processes of mineralization or solubilization of legacy P might not be sufficient for commercial crops, P is usually supplemented through fertilization, mostly as soluble phosphate (Solangi et al., 2023). However, being an anion, soluble phosphate added as fertilizer is rapidly fixed by positively charged Fe and Al oxides and hydroxides as well as by positively charged binding sites on organic matter and at the edges of phyllosilicates (Hinsinger, 2001). This can lead farmers to compensate through overfertilization, to assure a satisfactory level of available P. This behaviour has resulted in the global phosphorus cycle exceeding its planetary boundary, which is a proposed limit on the quantities of phosphorus fertilizers applied to soils (Blackwell, 2022). The low use efficiency of P by plants -either from fertilizers or from the soil legacy P- and the resulting overfertilization have led to increased accumulated P in soils that in turn can be lost through runoff or soil erosion. P in runoff and soil particles can end up in water bodies, where increased P can lead to eutrophication and excessive cyanobacterial growth (Bonilla et al., 2023) and derived environmental problems. This situation has been described as the phosphorus paradox, to account for the simultaneous scarcity and overabundance of this key nutrient (Loughfeed, 2011). In soybean, although P is required for processes like photosynthesis and biological nitrogen fixation, P fertilization has been associated with increased seed P concentration and reduced seed vigour (Krueger et al. 2013), a key indicator of seed quality. In turn, increased seed and grain P level -mostly in the form of phytate-is a negative factor in plant-based food and feed since phytate is considered as a anti nutritional factor in animals (Singer et al., 2023).

Therefore, technologies that improve the bioavailability of P to plants are needed to enhance utilization of legacy P from the soil, increase the use efficiency of P fertilizers and thus reduce the P input into agricultural systems (Rowe et al., 2016). In turn, such technologies would have positive impacts on the sustainability of agriculture by reducing P losses to the environment.

The role of microorganisms in nutrient cycling and plant nutrition has been exploited by selection of microbial strains bearing traits associated with plant growth promotion like the production of plant growth regulators, the elicitation of induced systemic resistance (ISR) and biofertilization. In particular, strains of *Bacillus* are known to produce organic acids, phosphatases and phytases (Zhao et al., 2021). Excreted phosphatases can solubilize inorganic P or mineralize the rather labile forms of organic P like nucleotides and phospholipids in soils, while phytases are enzymes that sequentially remove phosphate groups from myo-inositol 1,2,3,4,5,6-hexakisphosphate (phytate), the main storage form of P in plants and organic P in soils (Singh et al., 2020). Therefore, phosphatase and phytase activities of bacteria inhabiting the plant rhizosphere may contribute to their plant growth promoting effect (Idris et al., 2002). In 2016, the organic phosphorus workshop held in UK acknowledged the role of microorganisms in P cycling and prioritized the need for greater understanding of the metagenomics and functional microbial genes involved in organic P turnover (George et al., 2018) while the need to prioritize field experiments was stated in the 2023 workshop in Chile. In this regard, 16S rRNA gene metabarcoding can be used to address the structure and function of bacterial communities in soils (Orwin et al 2018), where they contribute to cycling of nutrients while being affected by root exudates and crop management techniques like P fertilization (Qu et al., 2020; Ren et al 2023.)

Recently, strain ILBB592 of *Priestia megaterium* (formerly *Bacillus megaterium*) was selected at INIA Uruguay due to its demonstrated plant growth promotion activities *in vitro* and *in planta* (Torres et al., 2024). Among its main features, this strain showed P solubilization and mineralization *in vitro*, great capacity to form biofilms in response to soybean seed exudates, production of indole acetic acid and acc deaminase, and acted as a nodule enhancing bacteria in pot experiments (Torres et al., 2024). However, its effect on plant P content was not evident, as it was for other strains like *Bacillus pumilus* ILBB44. In addition, it is acknowledged that field trials are necessary to verify the potential benefits of plant growth promoting rhizobacteira (PGPR) and P biofertilizers. In view of this, we set up a field trial to assess the effect of co-inoculating soybean seeds with formulated prototypes of *P. megaterium* ILBB592 and *Bacillus pumilus* ILBB44 together with a commercial inoculant of *Bradyrhizobium elkanii* in use in Uruguay. We hypothesized that co-inoculation with either strain -ILBB44 or ILBB592-would improve plant P uptake, nodulation, vegetative and reproductive parameters of the local soybean cultivar INIA Genesis 5602 in field conditions. In addition, we analysed the impact of the co-inoculated bacteria and superphosphate fertilization on the rhizospheric bacterial community, to gain an initial understanding of the ecological effects of these inputs on the structure and functions of the rhizospheric and bulk soils bacterial communities.

## Methods

### Biological materials

*Bacillus pumilus* strain ILBB44 and *Priestia megaterium* strain ILBB592, known to promote nodulation and P nutrition of soybeans (Torres et al., 2024) were formulated as dry powder by Calister (Lallemand Uruguay) at a concentration of 10^9^ spores/g and used in co-inoculation bioassays with a commercial rhizobial inoculant (Active N ®, Lage & Cia.) containing two strains of *Bradyrizhobium elkanii* (U-1301 and U-1302) on soybean seeds cv. INIA Genesis 5602.

### Seed treatment

Batches of two kg of seeds were placed in plastic bags and first treated with fungicide Fluidox ultra TBZ (ai fludioxonil, metalaxil, tiabendazol) (1 mL per kg of seeds) and insecticide Tiametox 350 FS (ai thiametoxan) (1.2 mL per kg of seeds). One hour after chemical application, seeds were inoculated with suspensions containing a mix of *B. elkanii* strains and either ILBB592 or ILBB44. To obtain these mixtures, 8 mL of Active-N and 4 mL of Add It® adherent were mixed and supplemented with 1 g of the dry powder formulations of ILBB44 and ILBB592. The final mixtures were added to the plastic bags containing 2 kg of chemically treated seeds and mixed thoroughly. Control treatments comprised the chemically treated seeds inoculated only with a mix of 8 ml Active-N and 4 mL of Add It adherent.

### Recovery of rhizobia and Bacillus from inoculated seeds

Bacterial counts on seeds were performed using a sub-sample of 90 co-inoculated seeds soaked in 250 ml Erlenmeyer flasks containing 90 ml sterile saline solution. After agitation for 15 minutes, plates containing a modified YEM medium (g.l^-1^: mannitol 10.0; K2HPO4 0.5; yeast extract 0.5; MgSO4.7H2O, 0.2; NaCl 0.1, and Congo Red 0.04) (Vincent, 1970), supplemented with 0.5 mM of SO4Mn and FeCl3.6H2O 0.01 g l^-1^. After agitation for 15 minutes, plates containing a modified YEM medium (g.l^-1^: mannitol 10.0; K2HPO4 0.5; yeast extract 0.5; MgSO4.7H2O, 0.2; NaCl 0.1, and Congo Red 0.04) (Vincent, 1970), were supplemented with 0.5 mM of SO4Mn and FeCl3.6H2O 0.01 g l^-1^. Colonies of *P. megaterium* and *B. pumilus* were visible after 48 hs and had a pinkish colour whereas *B. elkanii* colonies were translucent and visible after 7 days (Table 1).

**Table 1.** Bacterial recovered from soybean seeds after inoculation.

| Treatment | Strain | CFU seed<sup>-1</sup> |
| --- | --- | --- |
| **Control** | *Bradyrhizobium elkanii* (U-1301/U-1302) | 5.40 x 10<sup>6</sup> |
| **ILBB592** | *Bradyrhizobium elkanii* (U-1301/U-1302) | 4.33 x 10<sup>6</sup> |
|  | *Prestia megaterium* ILBB592 | 3.12 x 10<sup>5</sup> |
| **ILBB44** | *Bradyrhizobium elkanii* (U-1301/U-1302) | 5.70 x 10<sup>6</sup> |
|  | *Bacillus pumilus* ILBB44 | 3.17 x 10<sup>5</sup> |

**Table 2.** Chemical analysis of soil.

| Chemical analysis | Soil |
| --- | --- |
| pH (H<sub>2</sub>O) | 5.9 |
| N (%) | 0.23 |
| Organic C (%) | 2.73 |
| Bray I (µg P/g) | 7.7 |
| Nitrate N (µg N/g) | 4.2 |
| Ammonium N (µg N/g) | 4.6 |
| Ca (mEq/100g) | 12.2 |
| Mg (mEq/100g) | 3.5 |
| K (mEq/100g) | 0.89 |
| Na (mEq/100g) | 0.36 |
| Titrateable acid (mEq/100g) | 5.1 |
| CECpH<sub>7</sub> (mEq/100g) | 22 |
| Total bases (mEq/100g) | 16.9 |
| % Sat (Bases) | 76.8 |

**Table 3.** Main effect of phosphorus fertilization and co-inoculation with *Bacillus pumilus* strain ILBB44 and *Priestia megaterium* strain ILBB592 on root nodulation, N, P and K uptake, shoot dry weight and yield of soybeans grown in plots in year 2020/2021.

| Nodule dry weight (g plant<sup>-1</sup>) |  |  |  | P uptake (g plant<sup>-1</sup>) |  |  |  |
| --- | --- | --- | --- | --- | --- | --- | --- |
|  | -P | +P | Average |  | -P | +P | Average |
| Control | 0.08 ± 0.02 | 0.12 ± 0.03 | 0.10 a | Control | 0.08 ± 0.02 | 0.09 ± 0.01 | 0.08 a |
| ILBB44 | 0.14 ± 0.04 | 0.18 ± 0.06 | 0.16 b | ILBB44 | 0.09 ± 0.02 | 0.11 ± 0.02 | 0.10 b |
| ILBB592 | 0.10 ± 0.03 | 0.17 ± 0.06 | 0.13 b | ILBB592 | 0.11 ± 0.01 | 0.13 ± 0.02 | 0.12 c |
| Average | 0.11 a | 0.16 b |  | Average | 0.09 a | 0.11 b |  |
| K uptake (g plant<sup>-1</sup>) |  |  |  | N uptake (g plant<sup>-1</sup>) |  |  |  |
|  | -P | +P | Average |  | -P | +P | Average |
| Control | 0.59 ± 0.12 | 0.59 ± 0.06 | 0.59 a | Control | 1.25 ± 0.20 | 1.37 ± 0.12 | 1.31 a |
| ILBB44 | 0.79 ± 0.20 | 0.74 ± 0.12 | 0.77 b | ILBB44 | 1.52 ± 0.29 | 1.53 ± 0.21 | 1.53 ab |
| ILBB592 | 0.88 ± 0.08 | 0.77 ± 0.12 | 0.83 b | ILBB592 | 1.79 ± 0.24 | 1.73 ± 0.25 | 1.76 b |
| Average | 0.75 a | 0.70 a |  | Average | 1.52 a | 1.54 a |  |
| Shoot dry weight (g plant<sup>-1</sup>) |  |  |  | Yield (t ha<sup>-1</sup>) |  |  |  |
|  | -P | +P | Average |  | -P | +P | Average |
| Control | 29.33 ± 4.78 | 29.17 ± 2.32 | 29.25 a | Control | 3204.22 ± 307.01 | 3566.77 ± 177.37 | 3385.49 a |
| ILBB44 | 37.67 ± 7.19 | 31.34 ± 6.24 | 34.51 ab | ILBB44 | 3256.26 ± 499.93 | 4039.78 ± 418.39 | 3648.02 a |
| ILBB592 | 41.51 ± 3.29 | 36.81 ± 5.59 | 39.16 b | ILBB592 | 3273.14 ± 528.61 | 4129.20 ± 250.22 | 3701.17 a |
| Average | 36.17 a | 32.44 a |  | Average | 3244.54 a | 3911.92 b |  |

Mean values ± standard deviation (SD) in each column, the same letters are not significantly different at P < 0.05 (LSD Fisher).

### Experimental site

The trial was set at the experimental station INIA La Estanzuela, Colonia, Uruguay (34°20ʹS, 57°41ʹO). Soil type was a Eutric Brunosol with low available phosphorus (Table 2). The experimental site had no history of *Bradyrhizobium* inoculation or soybean cropping and consequently the most probable number of *Bradyrhizobium* per gram of soil was zero.

**Table 2.** Main effect of phosphorus fertilization and co-inoculation with *Bacillus pumilus* strain ILBB44 and *Priestia megaterium* strain ILBB592 on root nodulation, N, P and K uptake, shoot dry weight and yield of soybeans grown in plots in year 2019/2020.

| Nodule dry weight (g plant<sup>-1</sup>) |  |  |  | P uptake (g plant<sup>-1</sup>) |  |  |  |
| --- | --- | --- | --- | --- | --- | --- | --- |
|  | -P | +P | Average |  | -P | +P | Average |
| Control | 0.15 ± 0.09 | 0.12 ± 0.05 | 0,138 a | Control | 0.035 ± 0.007 | 0.036 ± 0.005 | 0,036 a |
| ILBB44 | 0.26 ± 0.08 | 0.24 ± 0.11 | 0,251 b | ILBB44 | 0.039 ± 0.005 | 0.049 ± 0.011 | 0,044 ab |
| ILBB592 | 0.21 ± 0.09 | 0.23 ± 0.10 | 0,219 ab | ILBB592 | 0.044 ± 0.006 | 0.052 ± 0.009 | 0,048 b |
| Average | 0,206 a | 0,199 a |  | Average | 0,039 a | 0,046 a |  |
| K uptake (g plant<sup>-1</sup>) |  |  |  | N uptake (g plant<sup>-1</sup>) |  |  |  |
|  | -P | +P | Average |  | -P | +P | Average |
| Control | 0.29 ± 0.04 | 0.29 ± 0.05 | 0,294 a | Control | 0.69 ± 0.12 | 0.68 ± 0.11 | 0,684 a |
| ILBB44 | 0.31 ± 0.04 | 0.38 ± 0.06 | 0,347 ab | ILBB44 | 0.76 ± 0.10 | 0.87 ± 0.18 | 0,815 ab |
| ILBB592 | 0.34 ± 0.04 | 0.42 ± 0.08 | 0,381 b | ILBB592 | 0.82 ± 0.11 | 0.95 ± 0.21 | 0,885 b |
| Average | 0,317 a | 0,365 b |  | Average | 0,756 a | 0,833 a |  |
| Shoot dry weight (g plant<sup>-1</sup>) |  |  |  | Yield (t ha<sup>-1</sup>) |  |  |  |
|  | -P | +P | Average |  | -P | +P | Average |
| Control | 17.41 ± 2.29 | 17.63 ± 3.54 | 17,51 a | Control | 3171.56 ± 92.47 | 3103.47 ± 365.17 | 3137.55 a |
| ILBB44 | 18.46 ± 2.33 | 21.42 ± 3.87 | 19,94 ab | ILBB44 | 3266.04 ± 163.72 | 3220.19 ± 286.95 | 3243.12 a |
| ILBB592 | 20.81 ± 2.82 | 25.64 ± 4.89 | 23,23 b | ILBB592 | 3146.62 ± 198.04 | 3315.06 ± 201.53 | 3230.84 a |
| Average | 18,89 a | 21,56 a |  | Average | 3194.74 a | 3212.91 a |  |

Mean values ± standard deviation (SD) in each column, the same letters are not significantly different at P < 0.05 (LSD Fisher).

### Experimental design and field layout

In this study, a randomized complete block design was used, with four replications. The treatments were the combinations of two inoculation strategies and two fertilization levels: Soybean seeds inoculated with *Bradyrizhobium elkanii* (control) and Soybean seeds co-inoculated with *Bradyrizhobium elkanii* together with formulated spores of either *Priestia megaterium* ILBB592 or *Bacillus pumilus* ILBB44, in soils with and without superphosphate fertilization.

### Phosphate fertilization

Phosphorus fertilization was calculated to increase the level of available P in the fertilized plots from 7 to 15 ppm (P-Bray). For that, 150 kg of superphosphate 0-40/40-0 + 4S was evenly applied on the surface to achieve 60 kg P205 ha^-1^. After fertilization, soils from the fertilized plots were sampled to verify the achieved available P, which reached 14.7 ppm.

### Sowing and harvesting

Sowing of the control and treated seeds was done with a pneumatic seeding machine (Hartwich). The experimental design was BCA with 4 replicates. Each plot comprised six rows stretching five meters long and with 40 cm separation between rows. A density of 20 seeds per meter was used. Planting dates were November 14^th^ 2019 and November 19^th^ 2020. The harvest of the experimental plots was carried out with a Wintersteiger combine on 6^th^ May 2020 and May 14^th^ 2021. A total of 16 linear meters were collected per plot, after removal of 0.5 m from the edge of each plot. The fresh grain yield and yield corrected for humidity (13%) were obtained for each plot.

### Sampling and analyses

For soil analyses, 5 soil samples were collected from each plot at 0.0–0.15 m depth, mixed and a random subsample was used to determine soil chemical attributes before the beginning of field trial before sowing. Soil samples were collected from plots with and without phosphate fertilization thirty days after sowing to verify the effect of P application on the soil.

For plant nodulation and dry weight (DW), plants were collected at two growth stages: V6 (six unfolded trifoliate leaves) and R3.5 (early pod-fill stage) (Fehr et al. 1971). To evaluate nodulation, five plants at the V6 stage in each plot were uprooted and carefully washed with water so as not to detach the nodules. The soybean plants were cut at the cotyledonary nodes. Nodules were removed and placed in an oven at 65 ◦C until constant DW was obtained. Nodule DW was expressed as grams per plant (g/plant). To estimate the dry weight of the plants, five plants at the R3.5 stage in each plot, were dried in an oven at 65◦C until constant dry weight was obtained.

For plant nutrition, the following nutritional evaluations were performed: (a) P, N and K foliar concentration in g kg^-1^ of dry matter, was determined by collecting the middle third of 20 leaves of the main ear insertion in each experimental plot in the female flowering stage, according to the methodology described in Cantarella et al. (1997).

For metabarcoding analysis, bulk and rhizospheric soil samples were collected at the soybean V6 stage. Rhizosphere soil was recovered from 5 plants sampled from each of three plots, totalizing 5 g of roots per composite sample and 3 biological replicates per treatment. Composite samples were subjected to Stomacher treatment followed by centrifugation according to Schreiter *et al*. (2014). Rhizosphere pellets were stored at −20°C until TC-DNA extraction. TC-DNA was extracted from 0.5 g of frozen bulk soil and from 0.5 g of frozen rhizosphere pellet (wet weight) by harsh lysis using a FastPrep-24 bead-beating system and the FastDNA Spin Kit for Soil (MP Biomedicals, Santa Ana, CA, USA) following the manufacturer instructions. TC-DNA quality was checked by agarose gel electrophoresis. The TC-DNA was stored at −20°C. Total DNA was used as PCR template, and 16S amplicon libraries were generated using the PCR primers 515F (5′-GTGCCAGC MGCCGCGGTAA-3′) and 806R (5′-GGACTACHVGGGTWTCTAAT-3′) (Caporaso et al., 2011).

### Amplicon analysis

The raw reads were processed using the DADA2 v. 1.18 package (Callahan et al., 2016) in R Studio software v.4.0.3. The sequences were filtered and trimmed with the following parameters: Trunclen c(240,160), maxN = 0, maxEE = c(2,2), truncQ = 2, rm.phix = TRUE. The reads were merged, chimeric sequences removed, and taxonomy assigned to each merged sequence (amplicon sequence variant: ASV), and taxonomy assigned with assignTaxonomy based on the SILVA SSU database release 138.1 (Quast et al., 2013). Sequences affiliated to eukaryota, chloroplasts, mitochondria at the domain level were discarded. Alpha and beta diversity analysis of bacterial communities were carried out on rarefied ASV tables. The microeco package (Liu et al., 2021) was used to calculate the microbial alpha diversity index. Beta diversity was evaluated through a principal coordinate analysis (PCoA) using weighted Unifrac metric distance (Pérez-Jaramillo et al., 2017; García-Giraldo et al., 2022), with the entire filtered ASV table normalized using function cumNorm from the R package metagenomeSeq (Paulson et al., 2016), with the CSS (cumulative-sum-scaling) normalization method.

For functional analysis, changes in rhizospheric soil bacterial community functions related to P mobilization between −P and +P with and without co-inoculated bacteria were predicted using PICRUSt2 (Douglas et al., 2020). Kegg orthologs (Kos) involved in P-solubilization (ppx, ppa, gcd), organic P-mineralization (phoA, aphA, appA, phnP, phnX, phoD, phoX, phoN, phytase), P-starvation response regulation (phoB, phoP, phoR), P-uptake and transport (pit, phnC, phnD, phnE, pstA, pstB, pstC, pstS, ugpA, ugpB, ugpC, ugpE) were predicted. Predicted counts of genes were used to cluster the different conditions and were visualized through heatmaps using the pheatmap R package (Kolde, 2018) based on total counts of each gene and on the relative frequency of each gene in relation to the total number of predicted genes within each condition.

### Statistical analysis

Two-ways anova was used for in planta data sets to analyse interaction between P fertilization and co-inoculation, and then main effects of fertilization and co-inoculation were analysed by Fisher. Co-inoculation effects on alfa and beta diversity of rhizospheric and bulk soil bacterial community were analysed for each fertilization regime.

## Results

### Plant response to P fertilization and co-inoculation

Plant parameters were evaluated in two consecutive years and analysed on their own. There was not statistically significant interaction between phosphorus fertilization and co-inoculation of bacteria and hence main effects of these factors were considered. In the first year, plant P uptake was enhanced by co-inoculation with ILBB592 whereas P fertilization showed no effect. In the second year, co-inoculation with ILBB44 and ILBB592 as well as P fertilization had positive effects on plant P. Plant N and K followed the same trend as plant P in both years, when they were augmented by co-inoculation with ILBB592, but there was no main effect of P fertilization in the second year for these nutrients.

Regarding nodule dry weight, it was increased by co-inoculation in both years, with a higher effect of ILBB44 than ILBB592. Besides this consistent pattern, statistical significance was evidenced for ILBB44 in both years and ILBB592 on the second year. Phosphorus fertilization had a positive effect on nodule dry weight on the second year only. Shoot dry weight was positively affected by co-inoculation with ILBB592 on both years while P fertilization showed no effect on either year. Finally, yield showed no statistically significant effect of co-inoculation or phosphorus fertilization in the first year, whereas in the second year, yield average was clearly improved by co-inoculation and P fertilization, although this effect was statistically significant for P fertilization only.

### Alpha and Beta diversities

Alpha diversity was lower in bulk soils than in rhizospheric soil independently of soil fertilization regime and seed co-inoculation treatment. In soils without P fertilization, there was no further differentiation among rhizospheric soils from different seed co-inoculation treatments, whereas in soils with P fertilization, the co-inoculation with ILBB592 produced higher diversity than the co-inoculation with ILBB44, with rhizospheric soil from non co-inoculated seeds showing an intermediate degree of diversity, not statistically different from either of the co-inoculation treatments.

Beta diversity also showed that bulk soils had a bacterial community composition that differed from rhizospheric soils, independently of soil fertilization regime and seed co-inoculation treatment. In soils without P fertilization, the composition of rhizospheric soils seemed to be affected by seed co-inoculation treatment and this trend was more pronounced when P fertilization was applied since rhizospheric soils from seeds co-inoculated with either ILBB44 or ILBB592 showed discretely different communities from rhizospheric soils from seeds inoculated only with *Bradyrhizobium*.

When looking into class compositional variations with and without P fertilization (Fig. 3A), Alphaproteobacteria, Actinobacteria, Bacilli, Thermoleophilia and Verrucomicrobiae were the dominants bacterial class in the bulk soil, adding to 70% of the relative abundance, with the class Alphaproteobacteria and Thermoleophilia representing 40%. In the bulk soil no relevant changes were observed when comparing the -P and +P treatments. In the rhizospheric soil, the class Alphaproteobacteria, Actinobacteria and Gammaproteobacteria represent 85% of the relative abundance, and when assessing the effect of P fertilization, it was observed that the +P treatment significantly enriched Actinobacteria and Bacilli and decreased Alphaproteobacteria. But when seeds were also co-inoculated with ILBB44 and ILBB592, the opposite trend was observed, when relative abundances of Actinobacteria and Bacilli were lowered and Alphaproteobacteria was increased. Regarding variations in the relative frequency of genera, *Sphingomonas* and *Bacillus* were the most abundant in bulk soil with and without P fertilization (Fig. 3B), but in rhizospheric soils without P, their relative frequencies were reduced. In soils with P addition, the reduction was not verified for *Bacillus*, which remained at the same frequency as in bulk soil. When ILBB44 and ILBB592 were added, *Bacillus* remained similar to the levels in bulk soil.

**Figure 1.**
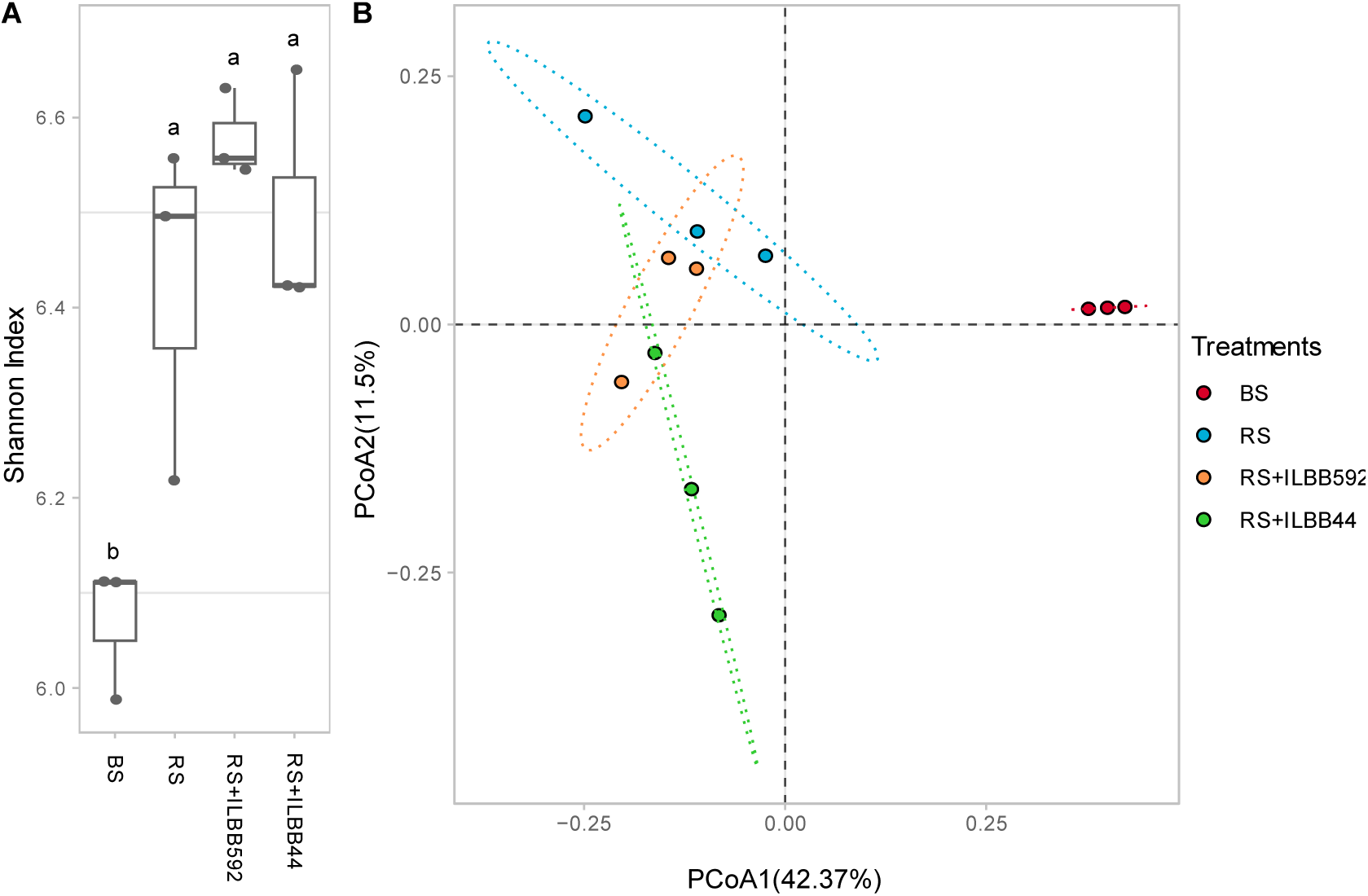
Principal coordinate analysis (PCoA) of bacterial diversity of soybean bulk soil and rhizospheric soil without P fertilizer. The treatments were: BS: bulk soil; RS: rhizospheric soil after seeds were inoculated with *Bradyrhizobium elkanii*; RS+ILBB592: rhizospheric soil after seeds were co-inoculated with ILBB592; RS+ILBB44: rhizospheric soil after seeds were co-inoculated with ILBB44. Data Was normalized with CSS and weight Unifrac metric distance was calculated.

**Figure 2.**
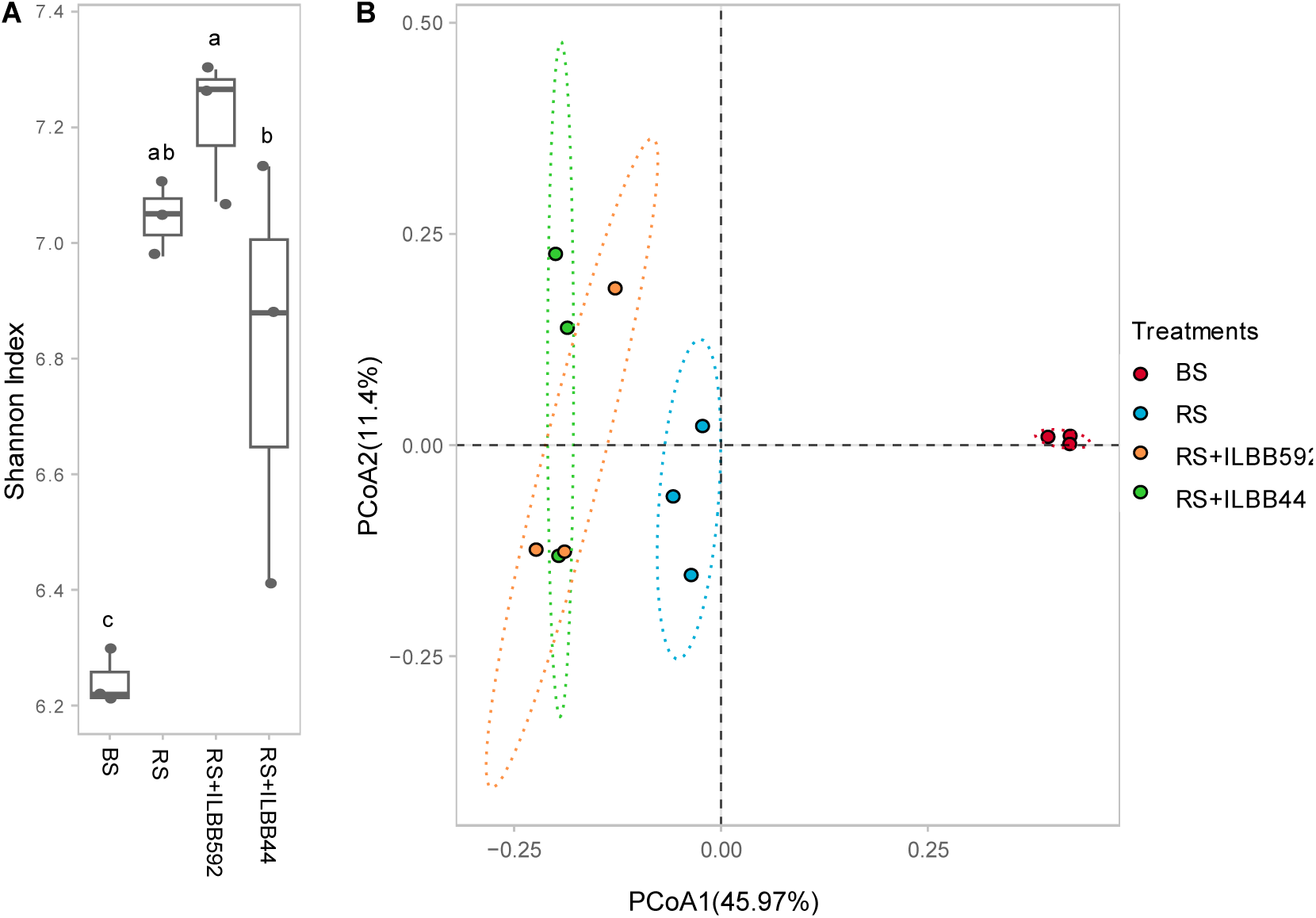
Principal coordinate analysis (PCoA) of bacterial diversity of soybean bulk soil and rhizospheric soil with P fertilization. The treatments were: BS: bulk soil; RS: rhizospheric soil after seeds were inoculated with *Bradyrhizobium elkanii*; RS+ILBB592: rhizospheric soil after seeds were co-inoculated with ILBB592; RS+ILBB44: rhizospheric soil after seeds were co-inoculated with ILBB44. Data Was normalized with CSS and weight Unifrac metric distance was calculated.

**Figure 3.**
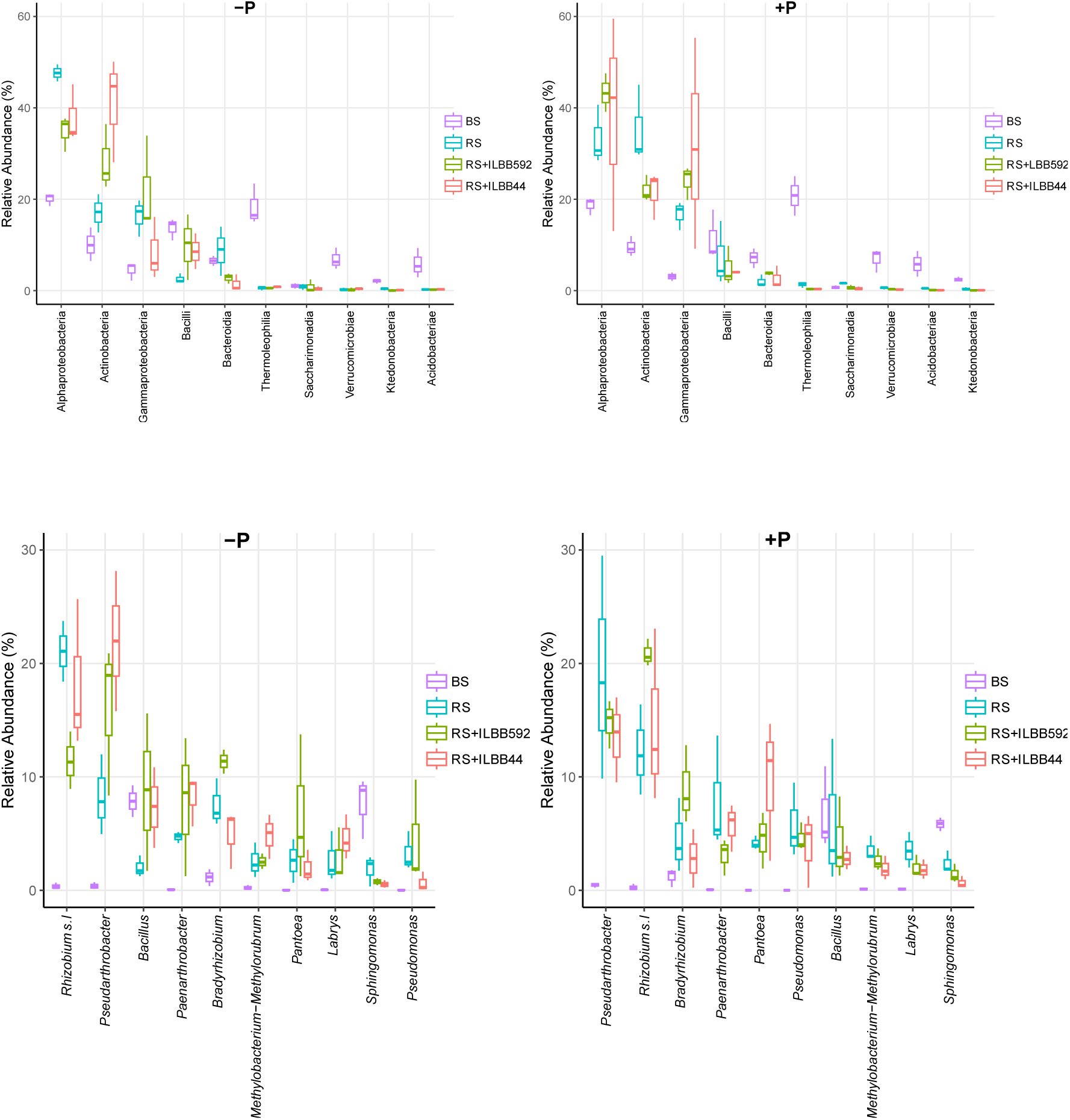
Effect of co-inoculation of soybean seeds on relative abundance of the most abundant Classes (A) and genera (B) in the non-fertilized (−P) and fertilized (+P) soils. BS: bulk soil; RS: rhizospheric soil after seed inoculation with *Bradyrhizobium elkanii*; RS+ILBB592: rhizospheric soil after seed co-inoculation with ILBB592; RS+ILBB44: rhizospheric soil after seed co-inoculation with ILBB44.

### Functional prediction

The number of genes predicted to be involved in inorganic P solubilization, organic P mineralization, P starvation response and P intake was higher in rhizospheric soil than in bulk soil, whereas the fertilization with superphosphate showed no effect on any gene category in either soil, except for predicted Inorganic P-solubilization genes, which seemed to decrease in bulk soil and rhizospheric soil when P fertilizer was added (Fig. 4). Within rhizospheric soils, the count of predicted genes in the four functional categories was highest when seeds had been co-inoculated with ILBB592, with and without P fertilization (Fig. 5). Predicted genes were lowest in the rhizospheric soil from seeds co-inoculated with ILBB44 without P fertilization while rhizospheric soil was the lowest when P was added, with P uptake and transport the most affected predicted gene category in the absence of P fertilization (Fig. 5).

**Figure 4.**
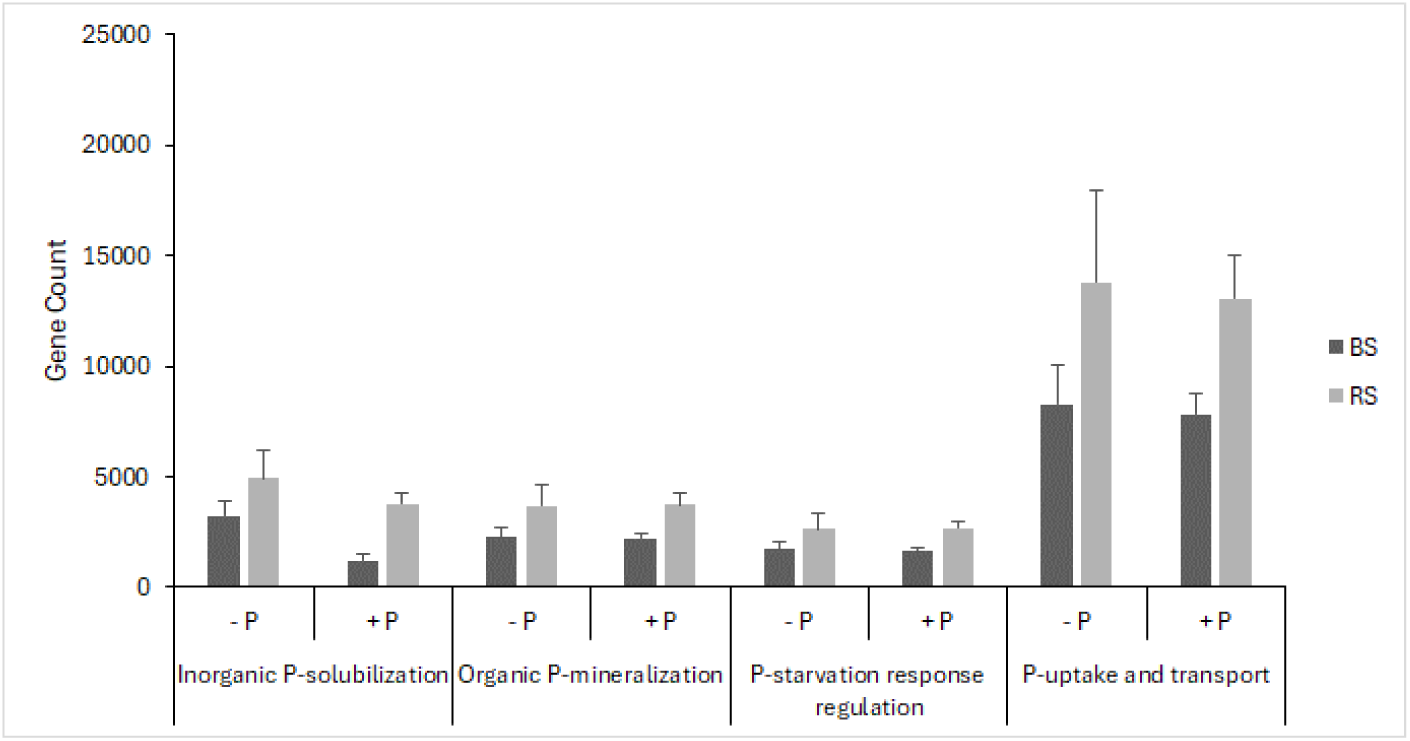
Abundance of genes within four gene categories involved in in inorganic P-solubilization (ppx, ppa, gcd), organic P-mineralization (phoA, aphA, appA, phnP, phnX, phoD, phoX, phoN, phytase), P-starvation response regulation (phoB, phoP, phoR), P-uptake and transport (pit, phnC, phnD, phnE, pstA, pstB, pstC, pstS, ugpA, ugpB, ugpC, ugpE) in rhizospheric soils from seeds inoculated only with rhizobia (RS) or bulk soil (BS) with and without superphosphate fertilization (+P/-P) at sowing.

**Figure 5.**
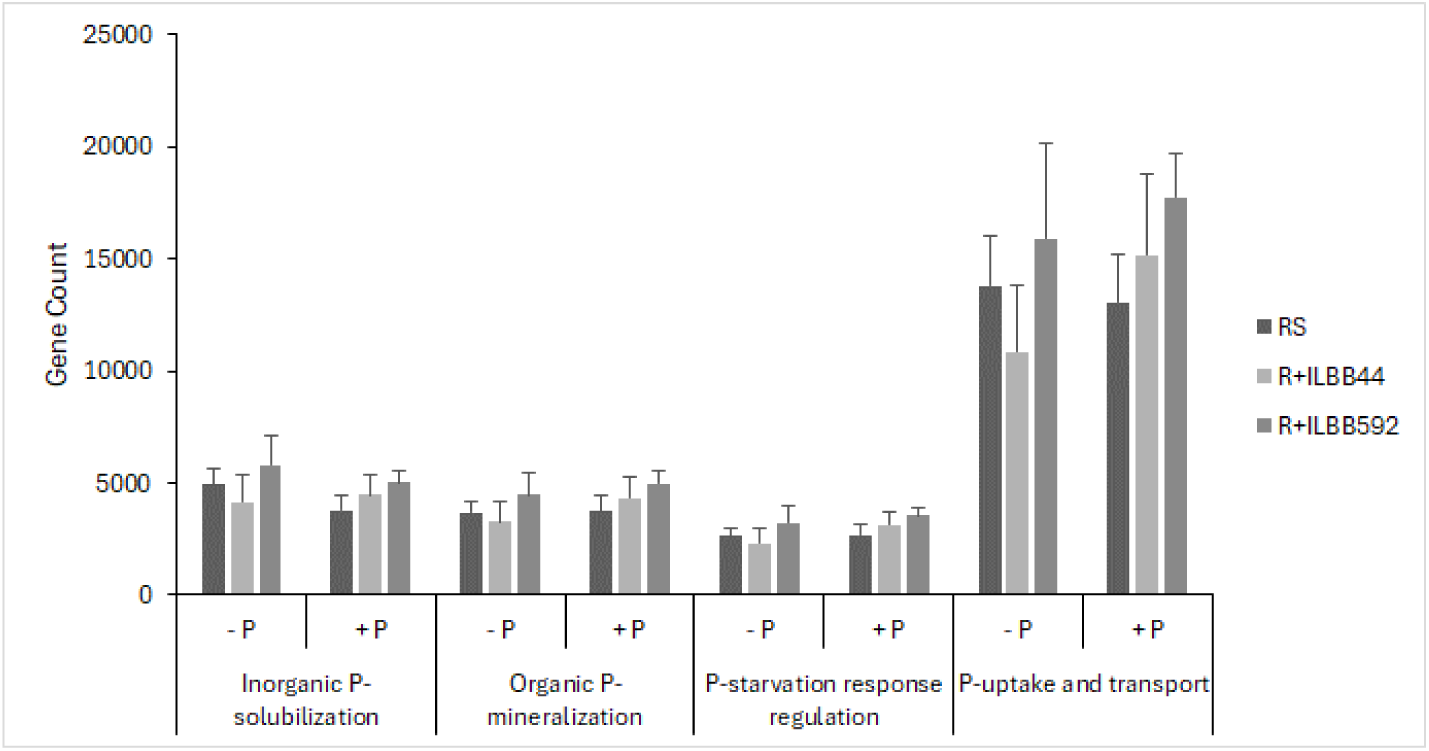
Abundance of genes within four gene categories involved in in inorganic P-solubilization (ppx, ppa, gcd, pqqA, pqqB, pqqC, pqqD), organic P-mineralization (phoA, aphA, appA, phnP, phnX, phoD, phoX, phoN, phytase, phoA), P-starvation response regulation (phoB, phoP, phoR), P-uptake and transport (pit, phnC, phnD, phnE, pstA, pstB, pstC, pstS, ugpA, ugpB, ugpC, ugpE) in rhizospheric soils from seeds inoculated only with rhizobia (RS) or co-inoculated with ILBB44 or ILBB592, with and without superphosphate fertilization (+P/-P) at sowing.

**Figure 6.**
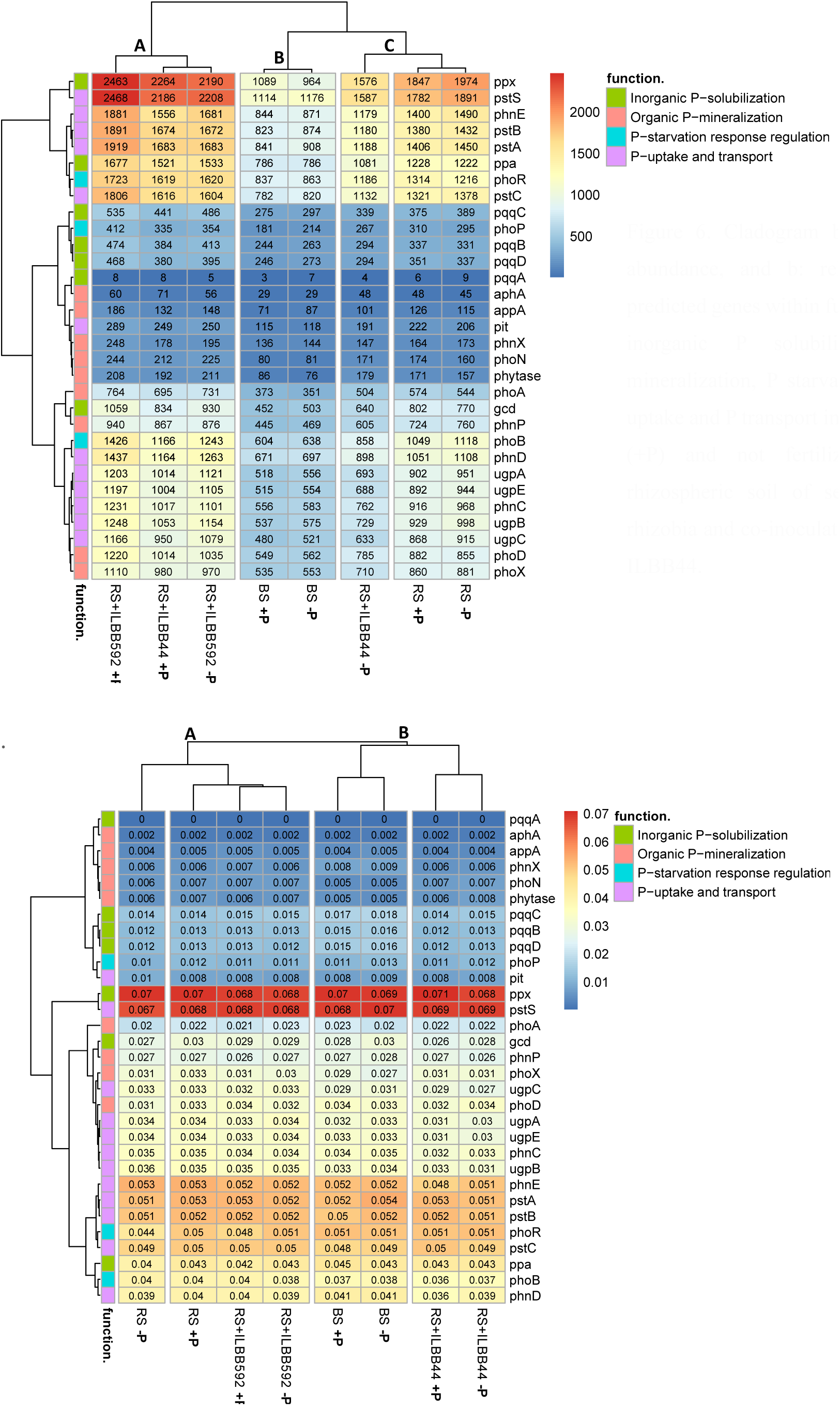
Cladogram based on a: absolute abundance, and b: relative abundance of predicted genes within functional categories of inorganic P solubilization, organic P mineralization, P starvation response and P-uptake and P transport in phosphorus fertilized (+P) and not fertilized (-P) bulk and rhizospheric soil of seeds inoculated with rhizobia and co-inoculated with ILBB592 and ILBB44.

When analysing the absolute abundance of each predicted gene related to P cycling, three clades were formed, with clade A comprising rhizospheric soils from seeds co-inoculated with ILBB592 (with and without P addition) and ILBB44 (with P fertilization only). This clade accounted for the highest numbers of predicted genes in all four functional categories, and within this clade, P fertilization and co-inoculation with ILBB592 showed the highest abundances of predicted genes. Clade B was formed by bulk soil (with and without P fertilization), with the lowest numbers of predicted genes in the four functional categories related with P cycling, and clade C -with an intermediate count of predicted genes-formed by rhizospheric soils (with and without P fertilization) and rhizospheric soils from ILBB44 co-inoculated seeds (without P fertilization). The predicted most abundant genes related with inorganic P solubilization were ppx, ppa and gcd ranging from 2469 to 1059 in rhizospheric soil from ILBB592 co-inoculated seeds (with P fertilization). Organic P mineralization genes were also highest in this condition, although phoD and phoX were the only genes above 1000. When analysing the relative abundances of predicted genes, differences were narrower than when analysing the absolute abundances. Still, two main clades were observed: clade A comprising rhizospheric soils from seeds inoculated only with rhizobia and from seeds co-inoculated with ILBB592 (with and without P fertilization), and clade B formed by two subclades, one comprising bulk soil and other comprising rhizospheric soil from seeds co-inoculated with ILBB44.

## Discussion

The role of *Bacillus sensu lato* in the bioavailablity of P has been studied and their development as biofertilizers has been proposed (Mosela et al., 2022; Vitorino et al. 2024). Although promising results have been obtained, there persist a need to further demonstrate activity in field situations for different crops. Because of this, we selected and formulated two strains, ILBB592 belonging to *Priestia megaterium* and ILBB44 ascribed to *Bacillus pumilus*, which had shown plant growth promotion and P mobilization potential *in vitro* and *in planta* in greenhouse assays (Torres et al. 2024), to assess their impact on P uptake by soybean cultivar INIA Genesis 5602. To assess the interaction of the selected strains with P added as fertilizer, we set up consecutive field experiments in plots with insufficient P Bray (7ppm) and plots supplemented with superphosphate to achieve 14,7 ppm P Bray, just above the critical P level of 14.2 ppm set for soybean yield in pampean soils in Argentina (Sucunza et al., 2018) and the mean P Bray in soils of 14 ppm in Uruguay (Bordoli et al., 2012). In addition to plant parameters, we evaluated the effect of P fertilization and co-inoculation on rhizospheric and bulk soil bacterial communities, to gain an initial insight on soil structural and functional changes underlying the observed effect on plant parameters.

Because soybean is routinely inoculated with *Bradyrhizobium elkanii* in Uruguay, we first evaluated the impact of co-inoculation with *P. megaterium* ILBB592 and *B. pumilus* ILBB44 on rhizobial nodulation and N nutrition. Co-inoculation with *Bacillus pumilus* ILBB44 outperformed ILBB592 and seeds treated only with *Bradyrhizobium* in both years, but plant N content was highest in plots co-inoculated with ILBB592. The highest increase in nodulation by ILBB44 could have come with an increased energy cost for the plants, beyond an optimal balance, as already observed in supernodulation mutants (Zhong et al., 2024), something that was also further supported by the intermediate shoot dry weight obtained by this treatment. This result might also be a first indication of ILBB592 enhancing plant performance and growth and therefore N accumulation, which was further supported by the highest values of shoot dry weight achieved by these plants on both years. Enhanced nodulation by ILBB592 had been reported in pot experiments with sand:vermiculite substrate (Torres et al. 2024), so now field experiments have contextualized the actual implications of this positive effect.

Plant P uptake was also augmented by co-inoculation with ILBB592, followed by co-inoculation with ILBB44 and then plants inoculated only with *Bradyrhizobium*. This increase was verified on both years, in plots with and without P fertilization. P fertilization had a positive effect on P plant content on the second year only. The variable effect of P fertilization on plant parameters has already been observed, and has been assigned to different factors like initial immobilization of P that can eventually become available in next cropping seasons (Sucunza et al., 2018). Beyond this, in both years, it was observed that when ILBB44 and ILBB592 had been co-inoculated, there was a consistent positive effect of P fertilization on P plant content. Therefore, in our experimental conditions, there seemed to be a conditional effect of P fertilization, which depended on the presence of ILBB44 and mostly ILBB592. This observation led us to analyse the bacterial community in the first year, when the soil with low P content of 7 ppm was receiving treatments for the first time. Although biodiversity and community structural changes are of interest, it has been proposed that soil processes can be better understood by the emergent soil functions (Vogel et al., 2018). Therefore, we evaluated changes in the diversity and composition of the bacterial community and particularly on soils gene functions related to P cycling, to acknowledge the effect of the different seed treatments on the rhizosperic microbiome at the light of the already analysed effect on plant parameters. The Shannon diversity index showed that diversity was higher in the plots that had received fertilization, and within each P fertilization regime, bulk soils exhibited much lower diversity than rhizospheric soils, and among these latter soils, those from seeds co-inoculated with ILBB592 had the highest diversity in both years, although these differences were statistically significant only on the second year. When exploring changes in the relative composition of the bacterial community, it was evident that class or genera that were prevalent in the rhizospheric soils were different from those prevalent in bulk soils, like Alphaproteobacterial and Actinobacteria doubling their relative frequency in rhizospheric soils, while the opposite was verified for other groups like Thermoleophilia and Bacilli. One case of interest was the relative abundance of class Bacilli and genus *Bacillus*, which were more prominent in bulk soil than in rhizospheric soils in plots without P fertilization, except when seeds had been co-inoculated with ILBB44 or ILBB592, when their relative frequency increased to similar level as in bulk soils. Their low frequency in rhizospheric soil was apparently counterbalanced either by the co-inoculation of ILBB44 or ILBB592 in unfertilized soil or by the addition of P fertilizer. When soil functions were explored through the prediction of a set of genes related to P cycling, it could be observed that the rhizospheric soil from seeds co-inoculated with ILBB592 in plots with P fertilization had the highest numbers of predicted genes in all four functional categories. This could be a sign of higher P cycling activity in the rhizosphere of these plants, which were also the ones with the highest plant P concentration. This treatment clustered with rhizospheric soil from seeds inoculated with ILBB44 with P fertilization and ILBB592 without P fertilization, which were therefore characterized by higher counts of the P cycling genes. Interestingly, these three treatments produced the three highest concentration of plant P. Altogether, there seemed to be a high association between the total number of P cycling genes in the rhizosphere and the plant phosphorus content. Similarly, it has been observed that P-solubilizing bacteria contribute to the restoration of degraded soils through enhancement of P cycling genes (Liang et al., 2020), and the application of bacterial strain *Acinetobacter pittii* gp-1 significantly increased soil available P, enriching both inorganic and organic P cycling related gene such as PhoD, bbp, gcd, and pstS (He and Wan, 2021).

Bulk soils, with and without P fertilization, showed the lowest number of predicted genes related with P cycling, while rhizosperic soils from seeds inoculated only with *Bradyrhizobium* had intermediate levels. When the relative frequency of the predicted genes in relation to the total numbers of genes predicted in each situation was calculated, apparent differences among treatments were not so clear, although they were sufficient to cluster soils by co-inoculation treatment, independently of the fertilization regime. Interestingly, rhizospheric soils from ILBB592 co-inoculated seeds clustered next to rhizospheric soils from seeds inoculated only with *Bradyrhizobium*. This might mean that soil function regarding P cycling is similar in both soils, but probably more intense when seeds were co-inoculated with ILBB592. This realization could be interpreted as a very positive aftermath of co-inoculation with this strain, with low distortion of rhizospheric equilibria/functions.

## Conclusions

*Priestia megaterium* ILBB592 had a positive effect on plant P, N and K accumulation on both years, independently from the phosphorus fertilization regime, and therefore phosphorus uptake was increased by seed co-inoculation with this strain. Related soil functions assessed by the count of predicted P cycling genes suggests that a higher number of these genes in the rhizosphere of ILBB592 co-inoculated seeds could be associated with higher P cycling and the highest level of plant P. The effect of phosphorus, on the other hand, was rather variable, being affected by the year in which it was evaluated and favoured by the co-inoculation of seeds with both ILBB44 and ILBB592. Although the role of rhizospheric microbiological processes in P fertilization efficiency and P bioavailability was highlighted, their interconnection with physical and chemical processes that explains the positive impact of co-inoculation was not addressed and remains elusive.

## Acknowledgement

This research was funded by Agencia Nacional de Investigacion e Innovación (ANII, Proyecto ALI_1_2014_1_5046), Instituto Nacional de Investigación Agropecuaria (INIA Uruguay), Institut Pasteur de Montevideo, Calister, Lage and Lafoner companies. Support and collaboration from staff of the institutes and companies involved is acknowledged.

## Notes

### Competing Interest Statement

The authors have declared no competing interest.

